# Microglial depletion decreases Müller cell maturation and inner retinal vascular density

**DOI:** 10.1101/2024.05.16.594541

**Authors:** Nathaniel Rowthorn-Apel, Naveen Vridhachalam, Kip M. Connor, Gracia M. Bonilla, Ruslan Sadreyev, Charandeep Singh, Gopalan Gnanaguru

## Abstract

The neuroretinal vascular system is comprised of three interconnected layers. The initial superficial vascular plexus formation is guided by astrocytes around birth in mice. The formation of the deep and intermediate vascular plexuses occurs in the second postnatal week and is driven by Müller-cell-derived angiogenic signaling. Previously, we reported that microglia play an important role in regulating astrocyte density during superficial vascular plexus formation. Here, we investigated the role for microglia in regulating Müller-cell-dependent inner retinal vascular development. Results from this study show that microglia closely interact with Müller cells and the growing inner retinal vasculature. Depletion of microglia resulted in reduced inner retinal vascular layers densities and decreased Vegfa isoforms transcript levels. RNA-seq analysis further revealed that microglial depletion significantly reduced specific Müller cell maturation markers including glutamine synthetase, responsible for glutamine biosynthesis, which is needed for angiogenesis. Thus, our study reveals an important role for microglia in Müller cell maturation and inner retinal angiogenesis.

## Introduction

Development of an intricate vascular network is important for the growth and survival of highly energy-demanding neural tissues such as the retina (1–3). The retinal vascular system is comprised of the interconnected superficial, intermediate, and deep vascular plexuses (1, 3). The growth of this complex three-layered vascular network is tightly regulated by multicellular interactions at distinct retinal developmental stages (1, 2, 4). In mice, during the first postnatal week, endothelial tip cells that emerge through the optic nerve head follow the astrocyte matrices to form the superficial vascular layer (5–8). Starting from the second postnatal week, endothelial tip cells that arise from the superficial vascular network, penetrate into the retina by following the Müller-cell-derived cues to form the deep and intermediate vascular plexuses (1, 2, 9, 10).

Müller cells are the most abundant intrinsically-derived glial cell type, which span almost the entire thickness of the retina (11, 12). Müller cells regulate a wide-variety of retinal functions including the regulation of inner retinal vascular growth and maintenance of the blood neural barrier (9, 12, 13). The morphogenesis of Müller cells begins at the end of the first postnatal week and continues to elaborate in the second postnatal week ahead of the inner retinal vascular developmental process (11). Müller-cell-derived cues, such as vascular endothelial growth factor (Vegf), attract the endothelial tip cells to penetrate into the retina to form the inner retinal vascular layers (2, 9). Müller cell specific deletion of transcription factor hypoxia-inducible factor (Hif) 2 alpha or Hif responsive growth factor (Vegf) during development leads to poor inner retinal vascular growth (9). Even in adulthood, ablation of Müller cells in the retina results in vascular abnormalities and breakdown of the blood neural barrier (13). This suggests that Müller cells play an important role in regulating inner retinal vascular growth and maintaining vascular integrity.

Besides Müller cells, microglia, the resident immune cell type, also closely interact with the endothelial tip cells and regulate vascular density during the course of the inner retinal vascularization process (14, 15). In particular, microglia that are located around the endothelial tip cells during the formation of the inner retinal vascular layers, modulate vascular density through non-canonical Wnt signaling and Tgfb1 levels (14, 15). Deletion of microglial Wnt ligand transporter (Wntless) or microglia dependent Tgfb1 signaling during the growth of the inner retinal vasculature results in excessive and abnormal vascular patterning (14, 15). This implicates that the microglia-derived cues are necessary for defining the density of the inner retinal vascular networks.

During the superficial vascular developmental process, prior studies including ours reported that microglia engulf and eliminate dying astrocytes in part through complement activation and facilitate the formation of spatially organized astrocyte/superficial vascular networks (7, 16). Intercellular interactions between Müller cells and microglia have been shown to facilitate the clearance of apoptotic cells during retinal development (17). It remains unclear if microglial-Müller cell interactions also modulate the density or the development of the deep and intermediate vascular plexuses in the retina. To investigate this, we explored the regulatory role of microglial-Müller cell crosstalk during the development of the inner retinal vascular layers.

## Results

### Endothelial tip cells closely interact with Müller cells and microglia during the formation of the inner retinal vascular plexuses

Retinal angiogenesis occurs in phases (1, 2), with the first phase of superficial vascular network formation being mediated by astrocytes (1, 5, 6). The next phase is the formation of the deep and intermediate vascular networks which are mediated by Müller cells, ultimately leading to the development of the complex interconnect three layered vascular network (Fig. 1A) (1, 2, 9). We recently reported that during the formation of the superficial vascular network, microglia, through complement activation, facilitate the spatial structuring of astrocytes and vascular networks (7). Here, we sought to determine if microglia closely interact with Müller cells to facilitate the growth of the deep and intermediate vascular plexuses during the second phase of retinal angiogenesis. We first examined retinal flatmounts for vascular, Müller cell, and microglial interactions during inner retinal angiogenesis (Fig. 1 and 2).

**Figure 1.**
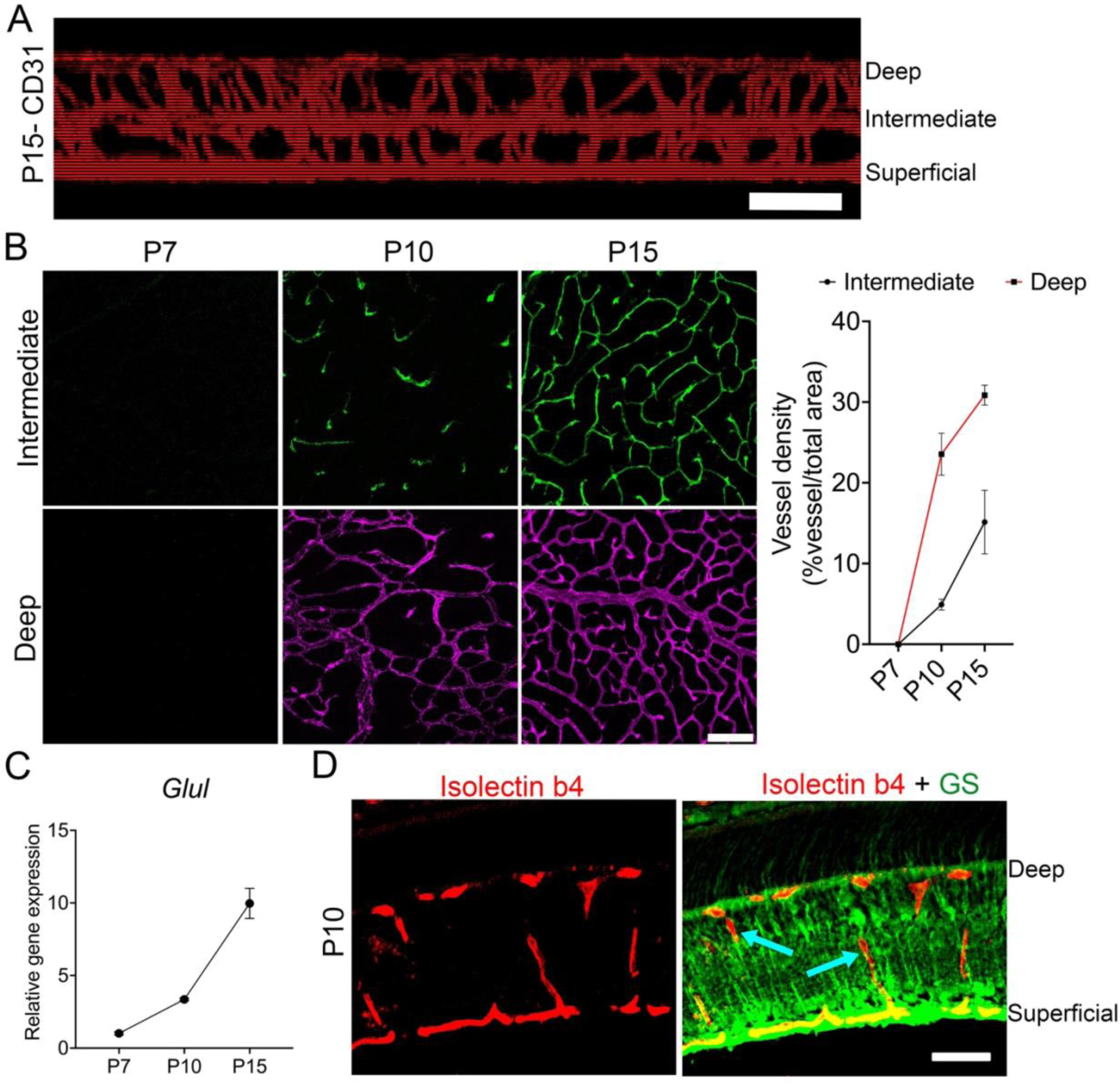
Endothelial cells migrate along Müller cell bodies to develop the inner retinal vascular layers. (A) Representative Z-projection of retinal flatmounts immunostained for CD31 revealing the interconnected superficial, intermediate, and deep vascular plexuses. (B) Retinal flatmounts immunostained for CD31 showing the vascular growth of the intermediate and deep vascular plexuses at P7 (n=3), P10 (n=4), and P15 (n=3). Graph quantifies vessel density age progression in the deep and intermediate vascular plexuses. (C) Graph shows relative mRNA expression of *Glul* at P7 (n=4), P10 (n=4), and P15 (n=5). (D) Representative P10 retinal cross section stained for Isolectin B4 (to label blood vessels) and glutamine synthetase (GS) (to label Müller cells). Arrows show endothelial tip cell migration over Müller cells. Scale bars: (A) is 20 µm, (B) and (D) are 50 µm.

**Figure 2.**
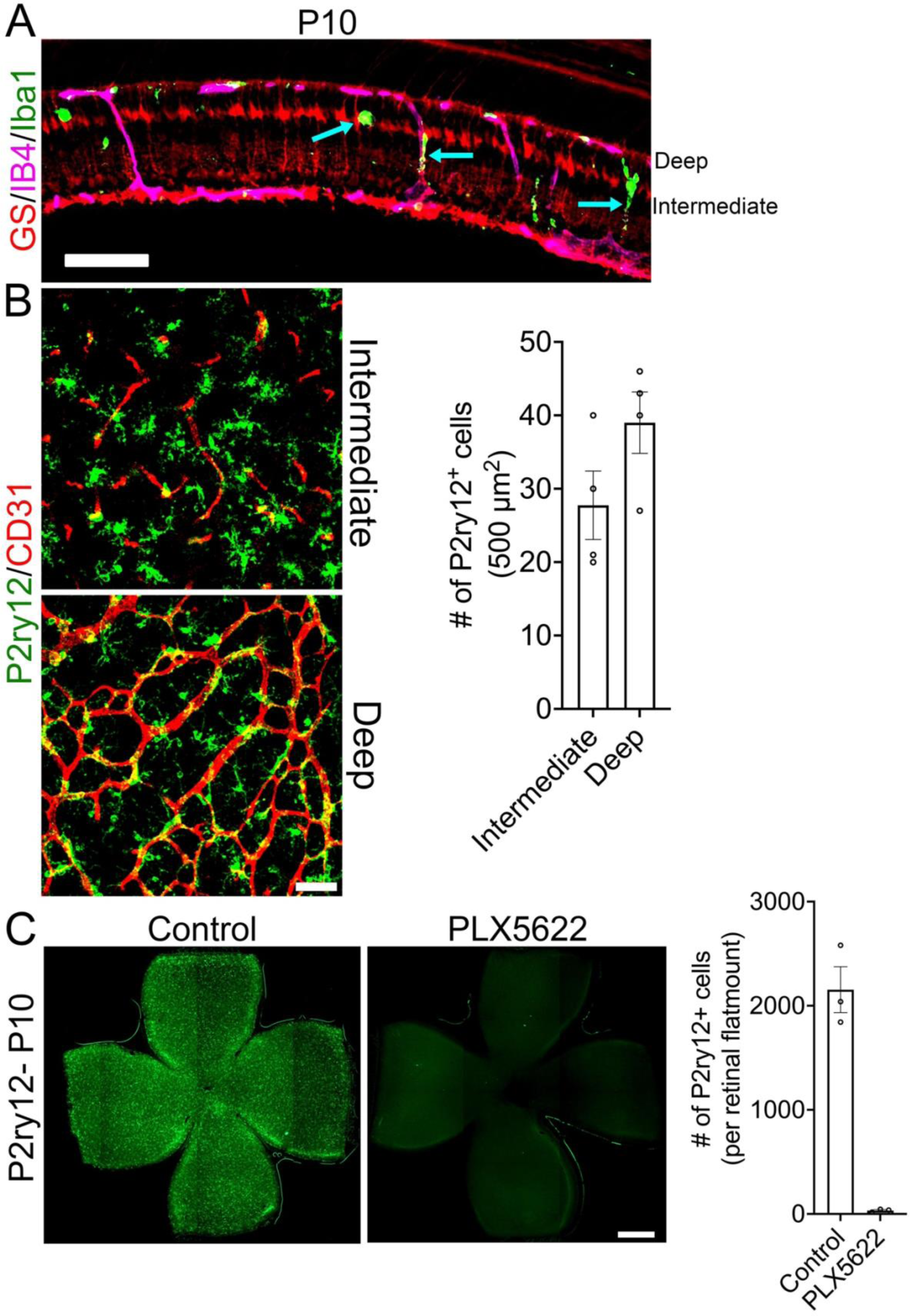
Microglia closely interact with endothelial cells and Müller cells during inner retinal vascular development. (A) Representative P10 retinal cross section immunostained for glutamine synthetase (GS), IsolectinB4 (IB4), and Iba1 revealing microglial association with the growing inner retinal blood vessels and Müller cells, (arrows). (B) Representative P10 retinal flatmount immunostained for P2ry12 and CD31 showing microglial association with the growing intermediate and deep vascular plexuses (n=4). Bar graph quantifies the number of P2ry12^+^ cells in the deep and intermediate vascular plexuses at P10 (n=4). (C) Representative P10 retinal flatmounts from control and PLX5622-treated groups showing microglial (P2ry12) distribution. Bar graph quantifies the number of P2ry12^+^ cells in the control and microglia depleted (PLX5622) groups and (n=3). Scale bars: (A) is 75 µm, (B) is 50 µm, and (C) is 500 µm.

Characterization of retinal flatmounts immunostained for CD31 shows that the growth of the inner retinal vascular plexus peaks at P10 and completes around P15 (Fig. 1A and 1B). Müller cells are the most abundant glial cell type of the retina that mature prior to inner retinal vascular plexus formation (Fig. 1C) (11) and drive the growth of the inner retinal vascular development (2, 9). Examination of P10 retinal crosssections revealed that the endothelial tip cells migrate along Müller cell bodies during the inner retinal vascular developmental process (Fig. 1D). In addition to Müller cells, microglia are also suggested to modulate inner retinal vascular density (14, 15). We therefore immunostained P10 retinal cross sections and flatmounts for a Müller cell marker (GS), microglial markers (Iba1 and P2ry12), and blood vessel markers (IB4 and CD31) to study microglial-Müller-endothelial interactions during the formation of the inner retinal vascular plexuses (Fig. 2A-B). Data revealed close association of microglia with the Müller cells and elaborating deep and intermediate vascular plexuses (Fig. 2A-B).

### Depletion of microglia reduces inner retinal vascular density

We next depleted microglia using a Csf1r antagonist (PLX5622) to study the effect of microglial loss on Müller cells and inner retinal vascular growth (Fig. 2C) (7). Following effective microglial depletion (Fig. 2C), we analyzed the CD31 immunostained retinal flatmounts of control and microglia depleted groups. Results show that the deep and intermediate vascular densities were significantly reduced by over 50% in microglia depleted retinas in comparison to the control retinas (Fig. 3A-B). Vegfa isoforms play an important role in driving the retinal angiogenesis (18, 19), we therefore determined the expression levels of Vegfa transcripts. Data show that all three Vegfa isoform (120, 164, and 188) transcript levels were significantly decreased in microglia depleted retinas in comparison to control retinas (Fig. 3C). Müller cells are the key source for Vegf in the retina that drive inner retinal vascular development and not the microglia (9, 20), we therefore performed mRNA sequencing on the P10 retinas of control and microglia depleted groups to determine the molecular link between microglial loss and reduced inner retinal vascular density.

**Figure 3.**
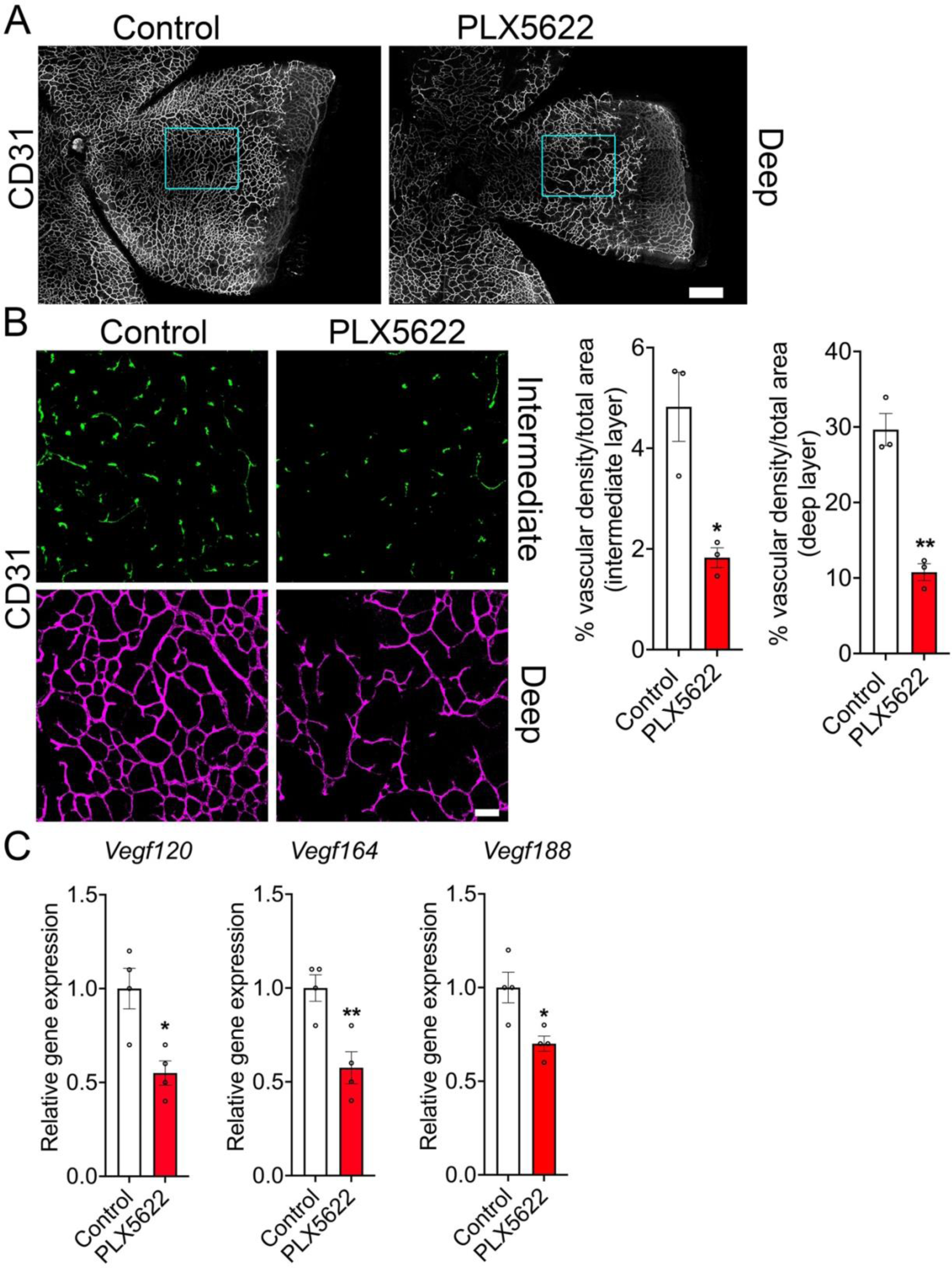
Depletion of microglia reduces vascular growth and *Vegfa* isoform expression levels. P10 control and microglia-depleted retinal flatmounts were immunostained for CD31 to examine deep and intermediate vascular plexus growth (A and B). (A) Retinal flatmounts of control and PLX5622 groups revealing the deep vascular plexus at P10. (B) Intermediate and deep vascular plexuses in the control and PLX5622 groups were imaged and quantified using ImageJ Angio Tool n=3 (boxed areas in A). (C) RT-PCR was performed to quantify the Vegf120, Vegf164, and Vegf188 isoform levels in the control and PLX5622 retinas at P10 (n=4). Scale bars: (A) is 200 µm and (B) is 50 µm. All error bars represent ± S.E.M. Statistical differences between control and PLX5622 group were calculated by an unpaired *t*-test. * P < 0.05, ** P < 0.01.

### Microglial depletion significantly reduced transcript levels of specific Müller cell markers

RNA-seq data further confirms that PLX5622 treatment significantly reduced the transcript levels of generic and signature microglial genes such as *P2ry12*, *Tmem119*, and *Fcrls* (Fig. 4A). In addition, consistent with previous findings, depletion of microglia led to increased expression levels of astrocyte markers such as *Pax2*, *Gfap*, *Pdgfra* (Fig. 4A) (7, 16). Intriguingly, we noticed significant downregulation of specific Müller cell markers such as *Glul*, Kir4.1 (*Kcnj10*), Kir2.1 (*Kcnj2*), *Aqp4* (Fig. 4A and supplementary Excel 1) (21). We then performed pathway analysis on the significantly differentially expressed genes using the Ingenuity Pathway Analysis© tool. The pathway analysis revealed that the genes involved in phagocytosis, complement system, and immune response were significantly downregulated in microglia depleted retinas compared to control retinas (Fig. 4B). Additionally, the pathway analysis tool revealed a significant downregulation of genes involved in water transport, potassium homeostasis, and glutamine biosynthesis (Fig. 4B), which are highly expressed by Müller cells (11, 21, 22).

**Figure 4.**
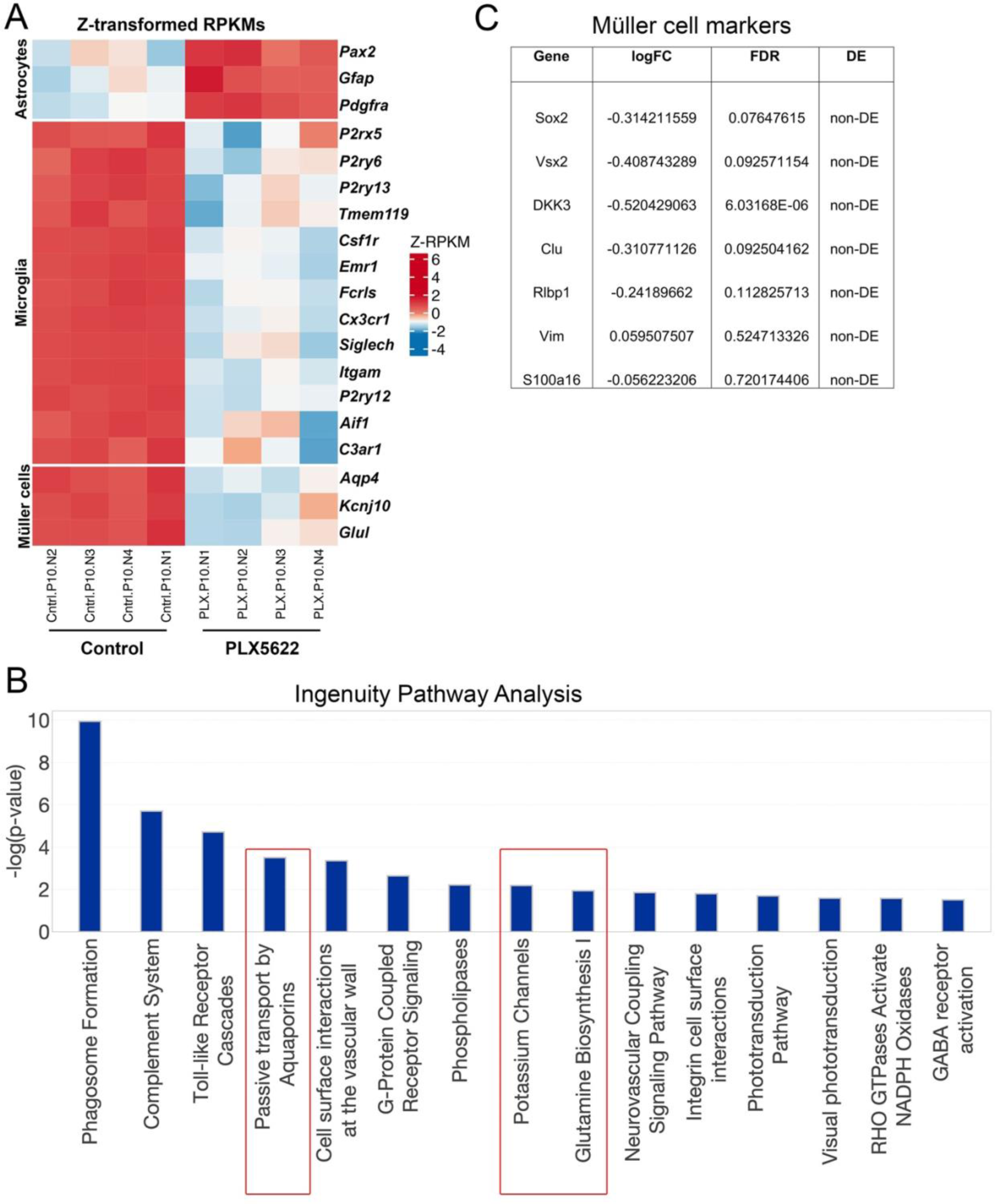
Microglial depletion significantly reduces Müller cell marker expressions. (A-C) Poly-A captured mRNA sequencing was performed on RNA extracted from the retinas of P10 control and PLX5622 treated groups (n=4). (A) Heatmap showing differential expression of significantly changed astrocyte, microglial, and Müller cell transcripts. (B) Ingenuity pathway analysis revealing top 15 significantly changed biological pathways in microglia depleted retinas in comparison to control retinas. (C) Table showing list of Müller cell specific transcripts that were not significantly changed in microglia depleted retinas compared to control retinas.

We examined for expression levels of other Müller cell markers in our RNA-seq data to determine if microglial depletion altered Müller cell growth. Our analysis did not show any significant differential expression of transcription factors such as Sox2 or Vsx2 expressed by Müller cells in microglia depleted retinas compared to control retinas (Fig. 4C) (23–26). Similarly, microglial depletion did not significantly alter the transcript levels of Sox8 and Sox9 transcription factors or its upstream regulator Hes5, which are involved in Müller development (supplementary excel dataset 1) (27). We also further examined the RNA-seq data and found no significant changes in the expression levels of Muller cell markers such as Dkk3, Clu, Rlbp1, Vim, or S100a16 in microglia depleted retinas compared to control retinas (Fig. 4C) (21). This suggest that microglial depletion alters specific gene expressional changes in Müller cells.

The downregulation of the glutamine biosynthetic pathway identified by the pathway analysis tool is of particular significance (Fig. 4B and 5A), because Müller cells are one of the major sources for glutamine release in the retina (28) and that glutamine is necessary for endothelial proliferation during angiogenesis (29, 30). To further validate the RNA-seq results, we used QPCR to measure and determine that the transcript levels of *Glul*, Kir4.1 (*Kcnj10*), and *Aqp4* were significantly downregulated in microglia depleted retinas compared to control retinas (Fig. 5B). Additionally, immunostaining of P10 retinal sections further confirmed that the depletion of microglia downregulated the localization of glutamine synthetase in Müller cells compared to controls (Fig. 5C). Of note, we did not notice significant differences in the transcript levels of glutamate receptors or transporters in microglia depleted P10 retinas compared to control retinas (Supplementary excel dataset 1).

**Figure 5.**
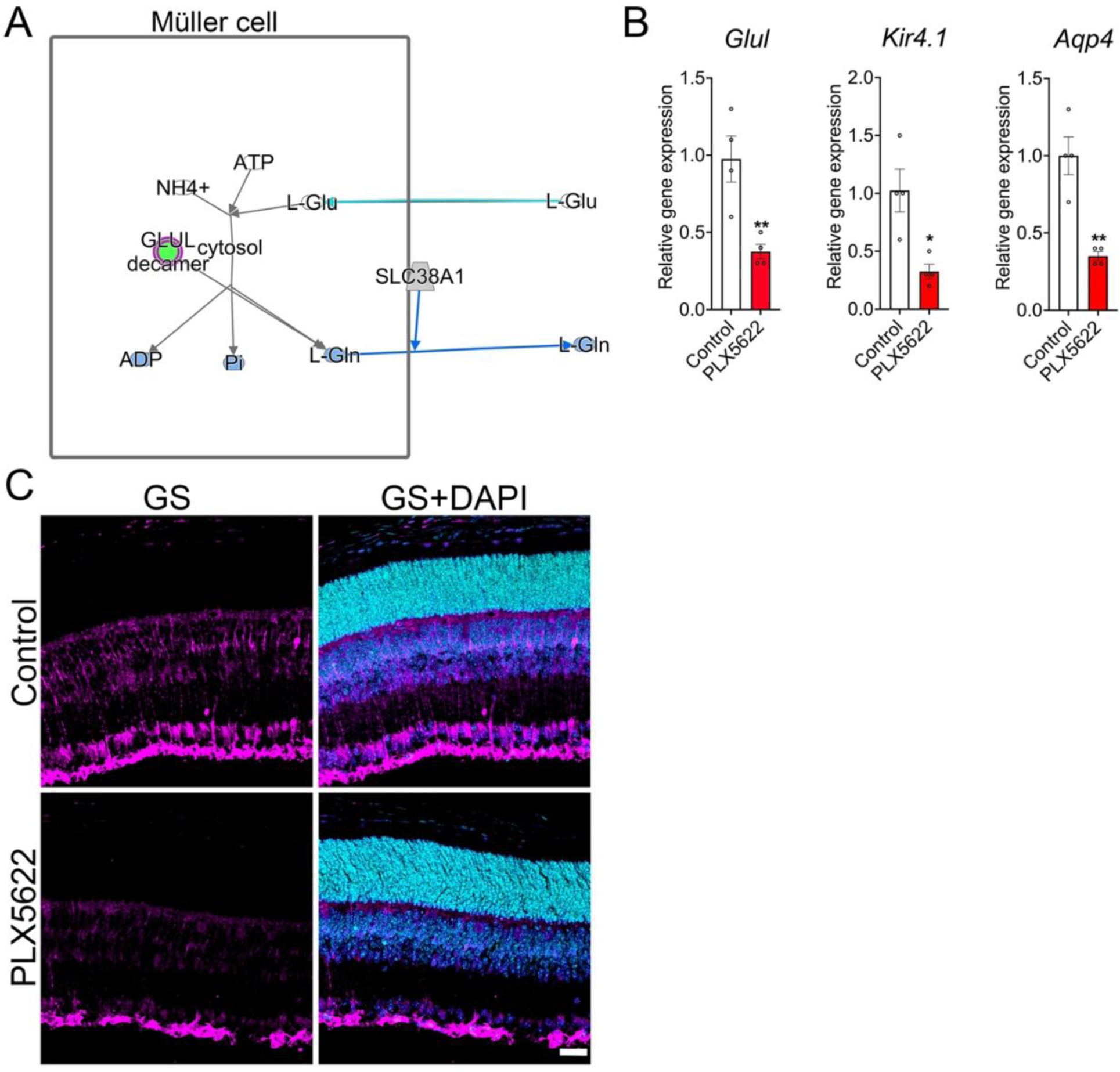
Microglial depletion results in significant downregulation of glutamine synthetase at P10. (A) Ingenuity pathway analysis showing significant downregulation of *Glul* gene (green) necessary for glutamine biogenesis and release in the microglia depleted retinas compared to control retinas at P10. (B) RT-PCR data revealing *Glul*, *Kcnj10* (Kir4.1), and *Aqp4* transcript expression levels between P10 control and microglia-depleted retinas (n=4). (C) Representative retinal cross sections showing immunostaining of glutamine synthetase and DAPI (nuclear stain) in control and microglia-depleted groups (n=3). Scale bar in (C) is 50 µm. All error bars represent ± S.E.M. Statistical differences between control and PLX5622 group were calculated by an unpaired *t*-test. * P < 0.05 and ** P < 0.01.

Thus, our data show that the depletion of microglia led to decreased deep and intermediate vascular plexus densities and diminished expression of specific Müller cell markers, suggesting an important role played by microglia during the second phase of inner retinal angiogenesis.

## Discussion

Microglia are a dynamic immune cell population of the retina that are distributed in distinct anatomical locations and regulate various developmental processes (7, 14, 31–34). Prior findings showed that the microglia closely associated with the astrocytes in the adjacent ganglion cell layer and eliminate the dying cells during the first postnatal week (7, 16), thereby facilitating spatially organized astrocyte template formation for the growth of the primary superficial vascular layer. However, the functional relevance of microglia closely interacting with the Müller cells in the inner retina, particularly during deep and intermediate vascular network formation, remains under characterized. In this study, we show that the microglia closely interact with the Müller cell body and the growing deep and intermediate vascular networks. Depletion of microglia led to a significant reduction in inner retinal vascular growth and decreased expression levels of Vegfa isoforms. Further RNA-seq studies revealed that the depletion of microglia resulted in diminished transcript levels of specific Müller cell markers that are necessary for cell maturation and blood vessel-retinal water exchange (35, 36).

Müller cell and microglial interactions were extensively characterized in the context of retinal homeostasis and injury models (37–40). The release of extracellular ATP, presumably by Müller cells, is suggested to modulate the morphology and the behavior of resting microglia (37, 39). Retinal injury models and co-culture systems show that the crosstalk between microglia and Müller cells elicits neurotrophic factor production and provides neuroprotection (38–41). Thus, prior investigations clearly reveal the importance of microglial-Müller cell interactions in retinal homeostasis (40). Recent findings show that microglial-Müller cell intercellular interactions are crucial for removing apoptotic cells during retinal development (17). The interaction between Müller cells and microglia during inner retinal vascular development is not clearly known. Immunostaining of retinal sections with Müller cell and microglial markers revealed close associations during active inner retinal angiogenesis. Hypoxia experienced in the developing inner retina activates Vegf expression in Müller cells, which drives endothelial tip cells to migrate into the retina (9). In addition to growth factor signaling, our data show that Müller cells potentially guide endothelial tip cells into the retina layers, similar to how astrocytes guide endothelial tip cells during superficial vascular development (42, 43).

During the migration of endothelial tip cells into the retinal layers in the second postnatal week in mice, microglia are shown to closely associate and regulate the inner retinal vascular density by suppressing excessive tip cell generation through non-canonical wnt signaling and by modulating Tgf beta signaling (14, 15). We therefore anticipated that the depletion of microglia would alter endothelial tip cell migratory patterns and/or increase inner retinal vascular density. In contrast to our expectation, microglia depleted retinas showed decreased densities of deep and intermediate vascular plexuses with significant reduction in the expression levels of Vegfa isoforms. RNA-seq studies further revealed that the depletion of microglia resulted in significant changes to selective Müller cell maturation markers. This suggest that the reduced inner retinal vascular growth observed in microglia depleted retinas is likely due to diminished expression of specific Müller cell maturation markers, in particular glutamine synthetase.

Glutamine metabolism plays an important role in retinal angiogenesis (29, 44). Depravation of glutamine impairs endothelial proliferation leading to reduced angiogenesis (29). Müller cells are the major source for glutamine biogenesis in the retina (28). Our pathway analysis revealed a significant decrease in glutamine biogenesis in microglia depleted retinas compared to control, suggesting that in addition to decreased Vegfa isoform levels, reduction in glutamine synthesis could also contribute to reduced deep and intermediate vascular plexus formation. It is still unclear how the depletion of microglia leads to downregulation of specific Müller cell marker expression such as *glul*. There are reports that suggest that the loss of specific cone photoreceptors, rod bipolar cells, and amacrine cells in the aged retinal degenerative (rd1) mouse model could lead to downregulation of glutamine synthetase reactivity in Müller cells_without changes in cell number (45). Although our RNA-seq studies did not reveal significant changes in the transcript levels of previously reported cell markers (45), there was significant decrease in bipolar cell markers such as Gpr179, a G-protein-coupled receptor required for mGluR6 signaling cascade in bipolar cells (supplementary excel dataset 1) (46). There was also significant decrease in Gnat1 (supplementary excel dataset 1), which is required for rods to rod bipolar cell synaptic transmission and rod photoreceptor survival (47). It is possible that the changes in bipolar and rod photoreceptor synaptic activity could contribute to down regulation of *glul* in Müller cells. Additionally, rod photoreceptors, Müller cells, and bipolar cells, originate from common progenitor cells (48), therefore, it is also possible that microglia directly or indirectly regulate certain genes expressions in the common retinal progenitors during maturation. Further studies are required to investigate if these developmental changes continue to persist in the adult retinas.

The vascular basement membrane is critical for the enrichment and sub-cellular localization of aquaporin and potassium channels in Müller cells (22, 49). The decrease in the expression levels of aquaporin and potassium channels observed in the microglia depleted retinas could be due to reduced vascular density.

Taken together, our data strongly implicates that microglia play an important role in facilitating Müller cell maturation, which is required for inner retinal vascular development. Depletion of microglia significantly reduces Müller cell maturation markers and inner retinal vascular density.

## Materials and Methods

### Mice and microglia depletion

C57BL6/J breeding pairs were purchased from Jackson Laboratories (stock # 00664, Bar Harbor, ME). Retinas were collected at P7, P10, and P15 for further analysis. For microglia depletion, timed-pregnant females at gestational day 13 or 14 were maintained on control chow diet (AIN-76A) or chow diet incorporated with Csf1r inhibitor (AIN-76A formulated at 1200 ppm) (PLX5622, Plexxikon Inc. Berkeley, CA). Retinas were collected from the control or microglia depleted (PLX5622) groups at P10 and used for downstream studies.

### RNA extraction and qPCR

Eyes enucleated and dissected in ice-cold 1x PBS to extract the retinas and then the RNA from the retinas were extracted using RNA-Stat-60 (Catalog# CS111, Tel Test Inc., Friendswood, TX) as per the manufacturer’s instructions. RNA concentrations were determined using the NanoDrop™ and the cDNA library was constructed using SuperScript™ IV VILO™ Master Mix (Catalog# 11756050, ThermoFisher Scientific, Waltham, MA). Real-time PCR reactions were performed in the CFX384™ Real-time PCR platform (Bio-Rad, Hercules, CA) using SYBR Green master mix (Applied biosystems, ThermoFischer Scientific, Foster City, CA) to determine the relative expression level of Kir4.1, Aqp4, Glul, Vegf120, Vegf164, and Vegf188. Sequences of primers used:

Ppia (TTCACCTTCCCAAAGACCAC, CAAACACAAACGGTTCCCAG),

Kir4.1 (GCCCCGTCTGTTCATCT, TGTAATAGACCTTAGCGACCGA),

Aqp4 (GGAAGGCATGAGTGACAGAG, TCCAGACTCCTTTGAAAGCC),

Glul (CATCCTGTTGCCATGTTTCG, CTCACCATGTCCATTATCCGTT),

Vegf120 (GCCAGCACATAGGAGAGATGAGC, CGGCTTGTCACATTTTTCTGG),

Vegf164 (GCCAGCACATAGGAGAGATGAGC, CAAGGCTCACAGTGATTTTCTGG),

Vegf188 (GCCAGCACATAGGAGAGATGAGC and AACAAGGCTCACAGTGAACGCT).

### Transcriptomic studies and pathway analysis

RNA was extracted as mentioned above from P10 control and microglia depleted retinas and the RNA integrity was examined using by Bioanalyzer (Agilent 2100). The RIN values of RNA used for cDNA library construction were >9.0. The RNA samples were treated with oligo d-T attached magnetic beads (NEBNext®) to pull down mRNA and used for cDNA library construction using NEBNext ultra II directional RNA library prep kit for Illumina as recommended by the manufacturer (New England Biolabs, Inc, Ipswich, MA). The quality of the cDNA libraries were assessed on the TapeStation (Agilent) and the libraries were sequenced using Illumina Nextseq 2000 platform with a target of 25million reads per sample.

For data analysis, STAR aligner was used for transcriptome mapping (50) and the mm9 assembly of the mouse reference genome. HTseq was used for obtaining individual Read counts (51) and the GENCODE M1 (NCBIM37) was used for gene annotation. EdgeR package was used for determining the differential expression of genes (52) after normalizing read counts and including only genes with CPM > 1 for at least one sample. Differentially expressed genes were defined based on the criteria of >1.5-fold change in normalized expression value was set to define the differential expression of genes and false discovery rate (FDR) <0.05. Heatmaps and PCA plots were generated using normalized gene expression values (log2 FPKM) and the Heatmaps were generated using the R package pheatmap (53). The pathway analysis of detected enriched genes were analyzed using Ingenuity® Pathway Analysis tool (Qiagen) version 01-23-01.

### Flatmount preparation and immunostaining

Flatmount preparations were performed as described previously (6, 7). Briefly, 4% Paraformaldehyde (BosterBio, Pleasanton, CA) fixed eyes (for 10 minutes), were dissected in 1x PBS and the retinal floatmounts were stored in ice-cold 100% methanol. Prior to staining, retinal flatmounts were washed in 1x PBS and then blocked (10% fetal bovine serum, 0.05%triton-X100, and 0.01% sodium azide in 1X PBS) for 2 hr at room temperature. Following blocking, retinal flatmounts were incubated with primary antibodies (see list below) for 24-36 hr at 4° C. After extensive washes in 1X-PBS, retinal flatmounts were incubated with respective secondary antibodies (see list below) for 4 hr at room temperature or overnight at 4° C. Following washes, retinal flatmounts were mounted onto slides using ProLong Gold anti-fade reagent (Invitrogen, Waltham, MA).

Primary antibodies used: goat anti-CD31 (1:500, Catalog # AF3628, R&D Systems, Inc. Minneapolis, MN). Isolectin b4 (1:250, Catalog#I21411, ThermoFisher Scientific, Waltham, MA), Glutamine synthetase (1:100, Catalog#AB305118, Abcam, Cambridge, United Kingdom), Rabbit anti-P2y12 (1:2000, Catalog# 69766, Cell Signaling Technology, Danvers, MA), and Goat Anti-Iba1 (1:500, Catalog# NB100-1028, Novus Biologicals, Centennial, CO).

Secondary antibodies used (1:500, all from ThermoFisher Scientific, Waltham, MA).: Donkey anti-Rabbit 647 (Catalog # A31573), Donkey anti-Rabbit 594 (Catalog # A21207), Donkey anti-Rabbit 488 (Catalog # A21206), Donkey anti-Goat 647 (Catalog # A21447), Donkey anti-Goat 594 (Catalog # A11058), Donkey anti-Goat 488 (Catalog # A11055).

### Cryosectioning

The eyes were enucleated and fixed in 4% PFA for 2hr at room temperature. Following fixation, the anterior chamber was dissected out and the posterior eyecups were cryopreserved for 24 hr in 30% sucrose. The eyecups were then embedded in Tissue-Tek® O.C.T compound (Ted Pella, Inc. Redding, CA) and frozen. Retinal sections (12µm thick) were made using cryostat and used for immunostaining studies.

### Imaging and analysis

The retinal sections and flatmounts were imaged using an epifluroscent microscope (Axio Observer Zeiss) or SP8, confocal microscope (Leica). The 3D-reconstruction of acquired Z-stack images were performed using NIH-ImageJ (version:2.1.0/1.53c).

Vascular density: Z-stack images of retinal flatmounts immunostained for CD31 were acquired (covering superficial till deep vascular plexus) using confocal microscopy. The vascular density in the intermediate and deep vascular plexuses were quantified using NIH-ImageJ software (version:2.1.0/1.53c) Angiotool plugin.

Quantifications of microglial cell numbers: Tiled images of retinal flatmounts immunostained for P2ry12 were acquired. Images were then analyzed using the NIH-ImageJ software (version:2.1.0/1.53c) by setting a threshold and quantifying the number of cells (determined by particle size) for each retinal flatmount and plotted.

### Statistical analysis

Prism version 9 software was used to perform all statistical analysis and the data were represented as the mean +/- SEM. Unpaired t-test was used to determine statistical differences experimental groups. In each figure, we noted the level of statistical significance and the ‘n’ for each experimental condition.

## Acknowledgments

This study was supported by NIH/National Eye Institute Grant R01EY032502 (to G.G) and NIH/NIDDK P30 DK040561 (to R.I.S). We thank Plexxikon, inc, for providing PLX5622 chow diet. We thank MGH Nextgen sequencing core staff Ulandt Kim for the help with cDNA library construction and sequencing.

## Author contributions

GG conceptualized, directed the study, and wrote the manuscript. KMC provided the resources and involved in experimental design. NR and NV performed the experiments and involved in manuscript writing. RS and GMB assisted with RNA-seq studies, and CS helped with data interpretation and manuscript preparation.

## Data availability

All the images and data analysis performed will be uploaded to the institutional repository (https://dataverse.harvard.edu/dataverse/tufts). The RNA-seq data will be upload to the GEO database repository. All the uploaded data will be made public upon acceptance and the data will also be made available by reaching out to the corresponding author: Gopalan Gnanaguru.

